# Inhibitory interneurons show early dysfunction in a SOD1 mouse model of amyotrophic lateral sclerosis

**DOI:** 10.1101/2020.10.21.348359

**Authors:** C. F. Cavarsan, P. R. Steele, L. T. Genry, E.J. Reedich, L. M. McCane, K. J. LaPre, A. C. Puritz, M. Manuel, N. Katenka, K. A. Quinlan

## Abstract

Few studies in amyotrophic lateral sclerosis (ALS) measure effects of the disease on inhibitory interneurons synapsing onto motoneurons (MNs). However, inhibitory interneurons could contribute to dysfunction, particularly if altered before MN neuropathology, and establish a long-term imbalance of inhibition / excitation. We directly assessed excitability and morphology of glycinergic (GlyT2 expressing) ventral lumbar interneurons from SOD1G93AGlyT2eGFP (SOD1) and wildtype GlyT2eGFP (WT) mice on postnatal days 6-10. Patch clamp revealed dampened excitability in SOD1 interneurons, including depolarized persistent inward currents (PICs), increased voltage and current threshold for firing action potentials, along with a marginal decrease in afterhyperpolarization (AHP) duration. Primary neurites of ventral SOD1 inhibitory interneurons were larger in volume and surface area than WT. GlyT2 interneurons were then divided into 3 subgroups based on location: (1) interneurons within 100 μm of the ventral white matter, where Renshaw cells (RCs) are located, (2) interneurons interspersed with MNs in lamina IX, and (3) interneurons in the intermediate ventral area including laminae VII and VIII. Ventral interneurons in the RC area were the most profoundly affected, exhibiting more depolarized PICs and larger primary neurites. Interneurons in lamina IX had depolarized PIC onset. In lamina VII-VIII, interneurons were least affected. In summary, inhibitory interneurons show very early region-specific perturbations poised to impact excitatory / inhibitory balance of MNs, modify motor output, and provide early biomarkers of ALS. Therapeutics like riluzole which universally reduce CNS excitability could exacerbate the inhibitory dysfunction described here.

**Abstract Figure:** Abstract Figure:
SOD1 glycinergic interneurons in the ventral horn show altered morphology and excitability, including depolarization of PICs, depolarized threshold, shorter AHPs, smaller somata and larger primary neurites. Ventrally located interneurons are the most prominently affected.

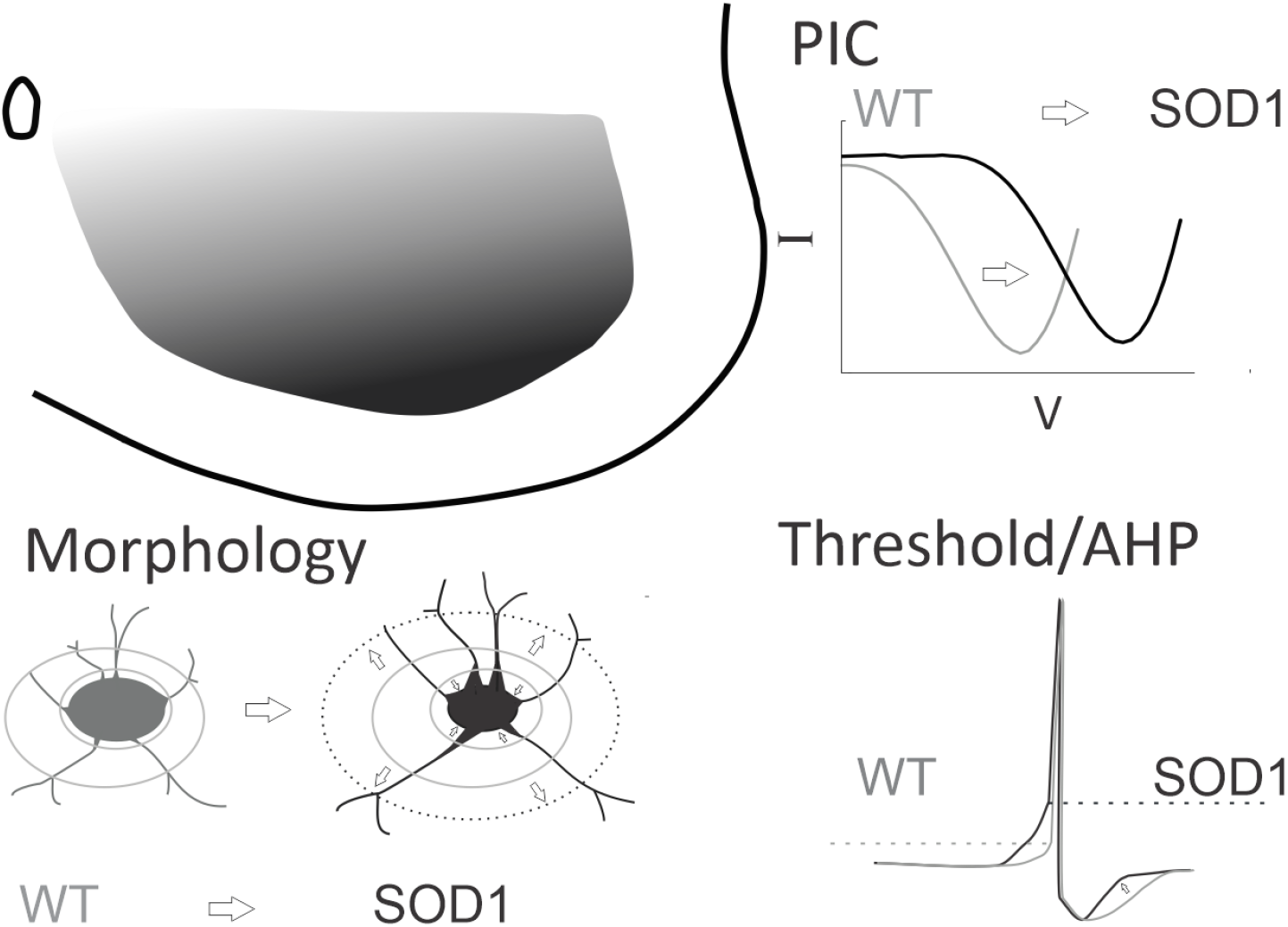

**Key Points Summary:**

- Spinal inhibitory interneurons could contribute to amyotrophic lateral sclerosis (ALS) pathology, but their excitability has never been directly measured.
- We studied the excitability and morphology of glycinergic interneurons in early postnatal transgenic mice (SOD1^G93A^GlyT2eGFP).
- Interneurons were less excitable and had marginally smaller somas but larger primary neurites in SOD1 mice.
- GlyT2 interneurons were analyzed according to their localization within the ventral spinal cord. Interestingly, the greatest differences were observed in the most ventrally-located interneurons.
- We conclude that inhibitory interneurons show presymptomatic changes that may contribute to excitatory / inhibitory imbalance in ALS.

## Introduction

Amyotrophic lateral sclerosis (ALS) is a rapidly evolving adult-onset neurological disease characterized by progressive loss of corticospinal neurons and spinal motoneurons (MNs). There has been considerable debate in the field over the role of hyperexcitability in neurodegenerative processes in ALS. While drug treatment for ALS based on nonspecific reduction of neuronal excitability (with riluzole, for example) (Bellingham, 2011), results in a modest increase in lifespan (Bensimon et al., 1994), similar treatment paradigms have been clinically disappointing. A more nuanced understanding of the excitability of vulnerable neurons could help to create a more targeted and effective approach for treatment of ALS. For example, if inhibitory pathways are failing in ALS and neuronal excitability is universally reduced with riluzole, this could further exacerbate inhibitory dysfunction by reducing activity not only in vulnerable MNs but also in inhibitory interneurons presynaptic to MNs. Thus, it is important to consider all aspects of neuronal excitability, including intrinsic excitability of vulnerable neurons, synaptic drive, and neuromodulation (Gunes et al., 2020). Intrinsic properties of MNs have been well studied but the same is not true for interneurons that are synaptically connected to them. In fact, no studies thus far have directly assessed activity of spinal premotor interneurons in an ALS model.

Despite evidence that ALS patients have disrupted inhibition at spinal levels (Raynor and Shefner, 1994; Shefner and Logigian, 1998; Sangari et al., 2016; Howells et al., 2020; Özyurt et al., 2020), much is still unknown concerning the involvement of inhibitory circuitry in ALS. Morphological alterations in inhibitory circuits have been demonstrated in animal models of ALS. These include degeneration of spinal interneurons, fewer neurons expressing markers of inhibitory neurotransmitters (including GlyT2) prior to MN loss (Martin et al., 2007; Hossaini et al., 2011), and loss of glycinergic boutons onto MNs prior to symptom onset (Chang and Martin, 2009a). Typically, fast MNs have greater numbers of inhibitory synaptic contacts, but these are largely lost in SOD1 mice beginning when motor unit atrophy is first observed (Pun et al., 2006; Hegedus et al., 2007; Allodi et al., 2021). Recurrent inhibitory circuits mediated by Renshaw cells (RCs) are impaired before symptom onset by both loss of MN collaterals which provide synaptic drive, and complex changes to RC-MN synaptic structures (Casas et al., 2013; Wootz et al., 2013). A few studies have suggested activity is decreased in inhibitory interneurons by indirect measurements. Quantification of synaptic inputs to MNs has shown that frequency of inhibitory postsynaptic potentials in spinal MNs is decreased embryonically in both SOD1 mouse and zebrafish models (McGown et al., 2013; Branchereau et al., 2019) and glycinergic inputs to MNs decay faster in SOD1 MNs (Medelin et al., 2016). On a larger scale, blocking the activity of V1 inhibitory interneurons during *in vivo* treadmill running was recently found to mimic the early locomotor deficits in SOD1 mice (Allodi et al., 2021), suggesting that inhibitory interneurons could be inactive or under-active in ALS. However, none of these studies directly examined electrophysiological activity in inhibitory interneurons.

We hypothesized that inhibitory spinal interneurons could contribute to the pathogenesis of ALS through a depression of MN inhibition. Decreased activity of inhibitory interneurons could result in synaptically-driven hyperexcitability of MNs and other long-term changes in network function. In this study, we examined electrical and morphological properties of glycinergic interneurons in the spinal cord of SOD^G93A^ GlyT2-eGFP (SOD1) mice compared to GlyT2-eGFP (WT) mice using whole cell patch clamp and three-dimensional reconstructions. We show here that significant dysfunction is present in SOD1 glycinergic interneurons, which likely comprise several subclasses of inhibitory interneurons. In general, SOD1 glycinergic interneurons are less excitable and have altered primary neurite morphology. Impairment in glycinergic interneurons should be explored as both a mechanism of vulnerability of MNs and a potential biomarker of early dysfunction of spinal circuits.

## Materials and Methods

### Ethical Approval

Experiments were performed in accordance with the United States National Institutes of Health Guide for Care and Use of Laboratory Animals and Northwestern University’s Animal Care and Use Committee. Approval was obtained for all experiments performed in this study (IS00001228), and all experiments conformed to the principles and regulations described in (Grundy, 2015). All efforts were made to minimize animal suffering and to reduce the number of animals used.

### Animals and Tissue harvest

Transgenic B6SJL mice overexpressing the human *SOD1^G93A^* gene (strain 002726, The Jackson Laboratory, Bar Harbor, ME, USA) and their wild type littermates (nontransgenic for the human *SOD1^G93A^* gene) were used. Transgenic animals were identified using standard PCR techniques by Transnetyx (Cordova, TN, USA) and were bred with GlyT2-eGFP mice obtained from Ole Kiehn’s lab at Karolinska Institute in Stockholm, Sweden, with permission of Hans Zeilhofer (Zeilhofer et al., 2005), generating SOD1^G93A^ GlyT2-eGFP mice, here called SOD1. Inhibitory glycinergic interneurons express the Na^+^ and Cl^-^ coupled glycine transporter 2, or GlyT2, and GlyT2-eGFP expression is driven by the GlyT2 promotor. Specifically, GlyT2-eGFP females were bred with SOD1^G93A^ GlyT2-eGFP males, and progeny were used for experiments before genotyping was performed. GlyT2-eGFP mice are referred to as wild type (WT), not carrying the SOD1^G93A^ mutation. All mice used in this study were housed at Northwestern’s Center for Comparative medicine and had *ad libitum* access to food and water. In total, 21 WT and 12 SOD1 juvenile mouse pups were used for the following studies. Mice were deeply anesthetized with isoflurane (Henry Schein Animal Health, Dublin, OH, USA) by inhalation until insensate to toe pinch, decapitated and eviscerated. The lumbar spinal cord from L1 - L6 was removed and embedded in 2.5% w/v agar (No. A-7002, Sigma-Aldrich, St Louis, MO, USA). The agar block was then superglued with Loctite 401 (Henkel Corporation, Rocky Hill, CN, USA) to a stainless-steel slicing chuck and 350 μm transverse slices were made using the Leica 1000 vibratome (Leica Microsystems, Buffalo Grove, IL, USA) as described previously (Quinlan et al., 2011). During both spinal cord isolation and slicing, the spinal cord was immersed in 1–4 °C high osmolarity dissecting solution containing (mM): sucrose 234.0, KCl 2.5, CaCl_2_ · 2H_2_O 0.1, MgSO_4_ · 7H_2_O 4.0, HEPES 15.0, glucose 11.0, and Na_2_PO_4_ 1.0. The pH was adjusted to 7.35 when bubbled with 95% O_2_/5% CO_2_ using 1 M KOH (Fluka Analytical, Sigma-Aldrich). After cutting, the slices were incubated for >1 h at 30 °C in incubating solution containing (mM): NaCl 126.0, KCl 2.5, CaCl_2_ · 2H_2_O 2.0, MgCl_2_ · 6H_2_O 2.0, NaHCO_3_ 26.0, glucose 10.0, pH 7.4 when bubbled with 95% O_2_/5% CO_2_ (all reagents for solutions were purchased from Sigma-Aldrich).

### Electrophysiology

Whole cell patch clamp was performed on interneurons from the lumbar segments using 2–4 MΩ glass electrodes pulled from glass capillary tubes (Item #TW150F-4, World Precision Instruments, Sarasota, FL, USA) with a Flaming-Brown P-97 (Sutter Instrument Company, Novato, CA, USA). Electrodes were positioned using a Sutter Instrument MP-285 motorized micromanipulator (Sutter Instrument Company). Whole-cell patch clamp measurements were performed at room temperature using the Multiclamp700B amplifier (Molecular Devices, Burlingame, CA, USA) and Winfluor software (University of Strathclyde, Glasgow, Scotland). Briefly, slices were perfused with a modified Ringer’s solution containing (in mM): 111 NaCl, 3.09 KCl, 25.0 NaHCO_3_, 1.10 KH_2_PO_4_, 1.26 MgSO_4_, 2.52 CaCl_2_, and 11.1 glucose. The solution was oxygenated with 95% O_2_/5% CO_2_, and the perfusion rate was 2.5 – 3.0 ml/min. Patch electrodes contained (in mM) 138 K-gluconate, 10 HEPES, 5 ATP-Mg, 0.3 GTP-Li and Texas Red dextran (150 μM, 3000 MW, from Invitrogen, Life Technologies, Grand Island, NY, USA). In voltage-clamp mode, fast and slow capacitance transients, as well as whole-cell capacitance, were compensated using the automatic capacitance compensation on the Multiclamp. Whole cell capacitance was recorded from the Multiclamp.

#### Neuron Selection

Glycinergic interneurons were visually selected for recording based on 1) expression of GFP, and 2) location in the ventral horn. Neurons that did not repetitively fire or did not maintain a resting membrane potential below −35 mV were excluded from electrophysiological analysis. Based on these criteria, 34/37 WT interneurons and 25/29 SOD1 interneurons were included in this analysis from mice between postnatal day (P) 6 - 10.

#### Electrophysiological analysis

Holding potential was set at −90 mV, and neurons were subjected to slow, depolarizing voltage ramps of 22.5 mV s^−1^, bringing the cell to 0 mV in 4 s, and then back to the holding potential in the following 4 s. In current clamp, neurons were subjected to depolarizing current ramps for testing I-on (the current level at firing onset), I-off (the current level at cessation of firing), and the slope of the frequency–current relationship. The difference between I-on and I-off is calculated for ΔI. Negative current was often necessary to prevent action potential (AP) firing. Resting membrane potential was recorded as the potential when 0 current was injected in voltage clamp. Persistent inward current (PIC) parameters were measured after subtraction of leak current. PIC onset was defined as the voltage at which the current began to deviate from the horizontal, leak-subtracted trace. PIC peak was the voltage at which the PIC reached peak amplitude. The linear portion of leak current (usually between −90 and −75 mV) was used to calculate whole cell input resistance. The first action potential in the train evoked by a depolarizing current ramp in current clamp mode was used to measure all parameters relating to action potentials. Threshold voltage was defined as the voltage at which the slope exceeded 10 V/s. Threshold for action potential firing was tested in two ways. The first was to use the voltage at firing onset from the current ramps up to 130 pA/s (in current clamp). The second was the voltage at which a single action potential could be evoked with a current step. Action potential overshoot is the voltage past 0 mV the first spike (from a ramp) reaches. Duration of the action potential is measured at half of action potential height (height is defined as overshoot – threshold voltage). Rates of rise and fall are defined as the peak and the trough of the first derivative of the action potential profile. Instantaneous firing frequency range max was the maximum firing rate that could be evoked from a neuron. It was measured from the first or second firing frequencies on a large (>150 pA) depolarizing step. Steady state firing frequency range minimum was the lowest firing frequency evoked from a small (^~^50 pA) depolarizing step. Hyperpolarizing steps were used to measure I_H_. The sag and rebound were measured from largest evoked response. Sag potential was calculated as a % of the total amplitude of the hyperpolarization. If a spike was evoked on rebound, the voltage threshold was used as amplitude of rebound. AHP duration was measured from a single spike fired during a period in which the membrane potential was stable. Amplitude was measured as the downward deflection from the membrane potential. Duration of the AHP was measured from the falling phase of the spike at baseline potential to the time of recovery to baseline membrane potential. Half amplitude duration was measured as the duration at half of the measured AHP amplitude from the falling phase of the spike at baseline potential. Tau is measured as the rate of decay from the last third of the AHP.

#### Regional classification

Regions were determined from spinal cord landmarks. In the case of patched neurons, photos were taken of the position of the patch electrode within the spinal cord slices after recording was complete. These photos were used to recreate a map of the position of the interneurons. For GFP+ neurons that were not patched but were reconstructed, z projections were used to group cells based on landmarks. The dorsal boundary of the ventral horn was identified by the central canal, the dorsal edge of lateral motoneuron pools and/or 300μm distance from the ventral white matter. Interneurons within the ventral-most region of the spinal cord (within 100 μm of the ventral white matter) were grouped into the RC region described previously (Siembab et al., 2010). Neurons that were located within and on the lateral edges of the motor pools (in between motoneurons and the white matter) were classified as lamina IX interneurons. Since the location of the motor pools varied throughout the lumbar enlargement, this classification was determined by examination of the z stacks and photos. Neurons that were not found within lamina IX and were not within the ventral RC region were grouped in the intermediate region of Lamina VII/VIII. All patched neurons were filled with Texas Red dextran to allow their 3D reconstruction. Reconstructions were limited to the first dendritic node (more details in following section). Unpatched GFP+ glycinergic interneurons that touched the boundary lines between zones were excluded from analysis to prevent double counting. When the position of GFP+ interneurons was not clear from the z stack, they were not included in the analysis.

### Morphological analysis

Neuron morphology was assessed from two types of images: 1) 2-photon image stacks of live neurons taken immediately after obtaining electrophysiological parameters, and 2) confocal image stacks taken after fixation and tissue processing. Images collected using these two methods were analyzed separately due to tissue shrinkage during the fixation process. Measurements from patched neurons included both soma and neurites (soma volume, soma surface area; and average neurite length [per cell], total neurite length [per cell], average and total neurite surface area and average and total neurite volume). Neurite reconstructions were performed up to the first node for several reasons, including variable dye filling of fine/distal processes and potential extension of neurites outside of the areas imaged. Neurites are modeled as the somata but by treating each branch as a frustum. Measurements of GFP+ glycinergic interneurons from fixed tissue samples included soma volume and soma surface area. Z-stacks acquired by confocal microscopy were used to generate 3D soma reconstructions in Neurolucida 360 (MBF Bioscience, Williston, VT, USA). Values shown throughout this study were calculated using Neurolucida’s branched structure analysis, specifically from the cell bodies summary page, including “enclosed volume” (calculated as the shell made from the cell body contours), and “surface area” (the sum of each section’s perimeter of the reconstruction multiplied by the thickness).

#### Two-photon imaging

An Olympus BX-51WIF microscope fitted with an Olympus 40x/0.8NA water-dipping objective lens was used. Two-photon excitation fluorescence microscopy was performed with a galvanometer-based Coherent Chameleon Ultra II laser (Coherent, Santa Clara, CA, USA) tuned to 900 nm. A red Bio-Rad 2100MPD photomultiplier tube (Bio-Rad, Hercules, CA, USA) (570 – 650 nm) was used to collect emission in the red channel. Z-stacks were obtained for each interneuron at 1024 × 1024 pixels (308 × 308 μm) resolution and roughly 100 μm in depth (step size 1 μm). From these Z-stacks, Texas Red® -filled neurons were three-dimensionally reconstructed, as described above, using Neurolucida software.

#### Confocal imaging

After patch clamp electrophysiology was complete, spinal cord sections were fixed, washed, and mounted with a coverslip for confocal microscopy. Z-stack photomicrographs (step size 1 μm) of glycinergic (GlyT2-eGFP+) interneurons in the lumbar ventral horn were imaged using a Nikon Eclipse Ti2 inverted confocal microscope at 20x magnification. From these Z-stacks, GlyT2 positive interneurons were three-dimensionally reconstructed using Neurolucida software as described above. The total number of analyzed interneurons was over 2000, including over 1000 from 6 WT mice and over 1000 from 6 SOD1 mice.

### Statistical analysis

Each patched neuron is treated as an independent observation. The assumptions of homogeneity of variances and the normality of the distribution of values for each measured characteristic were evaluated with Levine’s and Shapiro-Wilk tests, respectively. Group comparisons (WT vs SOD1) for patch clamp data were performed using one-way ANOVA on parameters that satisfied both conditions (homogeneity and normality of distribution) and using the Kruskal-Wallis test for those that did not. All results are presented as means +/- standard deviation (SD). Rather than basing our interpretation of results solely on *p* value, we have presented here the effect size of each measurement based on Hedges’ *g* value (Hedges, 1981). Effect size is presented using Cumming plots (Cumming, 2012) generated using DABEST, as in (Huh et al., 2021) for all patched neurons. Effect sizes are presented along with 95% confidence intervals (CI) based on a bootstrap distribution of the mean, resampled 5000 times and bias corrected. To compare anatomical data from unpatched neurons we used a random intercept linear mixed-effects model to account for the random effects of the mouse and the fixed effect of genotype; this was performed due to the correlation between data points created by the large number of repeated measurements from individual mice. The linear mixed model was used to avoid errors of pseudoreplication (Lazic, 2010; Yu et al., 2022) and dependency of *p* value on sample size (Lin et al., 2013; Gómez-de-Mariscal et al., 2021).

## Results

### Morphology of glycinergic interneurons

Ventrally located glycinergic interneurons in the lumbar enlargement of WT and SOD1 mice at postnatal day (P) 6-10 were reconstructed from z-stack images obtained using confocal microscopy. Typical ventrolateral glycinergic neurons are shown in **Figure 1**. Complete reconstructions of soma morphology were performed on over 2000 GlyT2 interneurons (1050 interneurons from 6 WT mice and 1170 interneurons from 6 SOD1 mice). Overall, soma size was not different: results of a linear mixed model did not indicate significance in either the soma surface area or enclosed volume, as shown in **Table 1**, and as can be appreciated by the lack of difference in distribution between the two populations as shown in Fig 1B. Analyzing dendritic morphology was not possible in these images due to the high number of GFP+ neurons and processes but was performed on a smaller subset of neurons that were imaged live after patch clamp electrophysiology with a 2-photon microscope.

**Figure 1:**
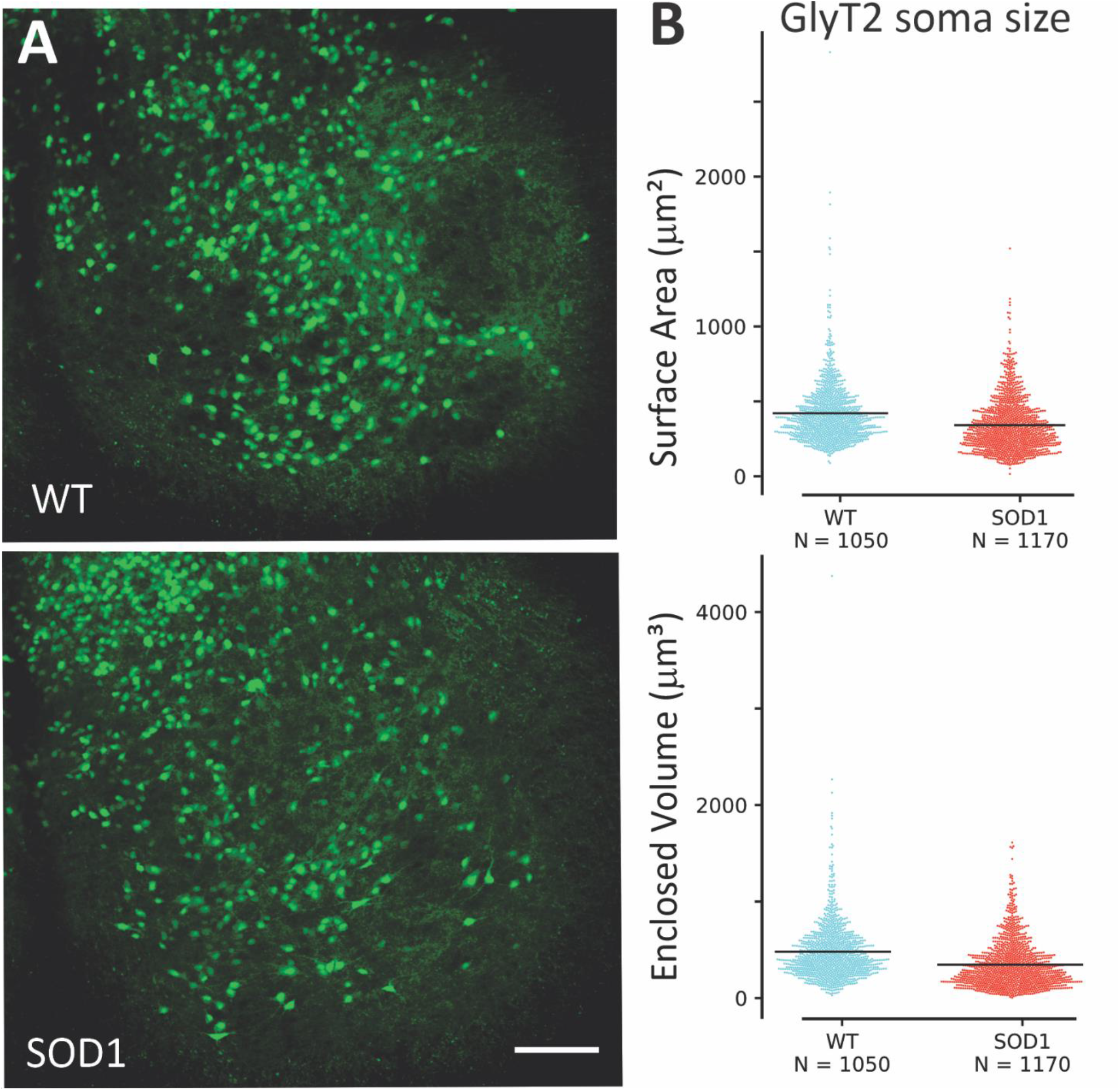
Comparing size of GlyT2 interneurons in the ventral horn. Representative photomicrographs of spinal cord showing GFP interneurons (green) in **A**. (**B**) Morphological analysis showed there was little difference in size between SOD1 glycinergic interneurons in ventral horn (N = 1170 neurons from 6 mice) and WT (N = 1050 neurons from 6 mice). Based on a linear mixed model differences are not significant. Scale bar in **A**: 100 μm applies to both images.

**Table 1:** Reconstruction data of GlyT2-GFP+ interneurons in WT and SOD1 animals.

| Ventral horn GlyT2-GFP+ Interneurons (somata only) |  |  |  |  |  |
| --- | --- | --- | --- | --- | --- |
| | | WT (Mean ± SD) | SOD (Mean ± SD) | *p* | |
|  |  | N = 1050 | N = 1170 |  |  |
| **Soma** | Surface Area ($\mu\text{m}^2$) | 450 (± 208) | 390 (± 178) | 0.387<sup>LM</sup> | |
| | Volume ($\mu\text{m}^3$) | 509 (± 306) | 414 (± 247) | 0.335<sup>LM</sup> | |
| Patched GlyT2-GFP+ Interneurons |  |  |  |  |  |
| | | WT (Mean ± SD) | SOD (Mean ± SD) | *p* | *g* (95% CI low, high) |
|  |  | N = 33 | N = 24 |  |  |
| **Soma** | Surface Area ($\mu\text{m}^2$) | 638 (± 232) | 653 (± 239) | 0.728<sup>K</sup> | 0.065 (-0.467, 0.621) |
| | Volume ($\mu\text{m}^3$) | 1472 (± 952) | 1643 (± 951) | 0.414<sup>K</sup> | 0.177 (-0.350, 0.731) |
| **Primary** |  |  |  |  |  |
| **Neurites** | Total Neurite Length ($\mu\text{m}$) | 101 (± 100) | 104 (± 49) | 0.905 | 0.029 (-0.441, 0.649) |
| | Average Neurite length ($\mu\text{m}$) | 29 (± 20) | 31 (± 13) | 0.663 | 0.109 (-0.387, 0.745) |
| | Total Surface Area ($\mu\text{m}^2$) | 513 (± 405) | 634 (± 266) | 0.032<sup>K</sup> | 0.338 (-0.230, 0.978) |
| | Average Surface Area ($\mu\text{m}^2$) | 152 (± 81) | 195 (± 81) | 0.025<sup>K</sup> | 0.523 (-0.035, 1.092) |
| | Total Volume ($\mu\text{m}^3$) | 265 (± 227) | 382 (± 190) | 0.009<sup>K</sup> | 0.546 (-0.023, 1.202) |
| | Average Volume ($\mu\text{m}^3$) | 79 (± 54) | 119 (± 61) | 0.008<sup>K</sup> | 0.679 (0.097, 1.245) |

N is the number of cells included in analysis; SD is standard deviation. Values of *p* are calculated from ANOVA except when marked with <sup>K</sup> indicating Kruskal-Wallis Test for nonparametric distributions or <sup>LM</sup> indicating linear mixed effects model. Values of *g* are determined using Hedge's statistic to calculate effect size from raw data and include the 95% confidence interval of the effect size

To study electrophysiological properties of GlyT2 interneurons, 59 glycinergic neurons (34 WT and 25 SOD1) were recorded using visually guided patch clamp with Texas Red dye in the electrode as shown in **Figure 2**. The location of patched neurons was distributed throughout the ventral horn as shown in Fig 2B. In this sample, we were able to analyze morphology of neurites up to the first node (branching point) and found that neurites were larger in patched SOD1 GlyT2 interneurons, including a modest increase in both neurite surface area and volume (see Table 1 for complete results). The number of primary neurites was not different in SOD1 vs WT interneurons. Please note that dimensions of these neurons cannot be directly compared to the previous section since these neurons were imaged in living tissue while the previous analysis was performed in fixed tissue, which shrinks.

**Figure 2:**
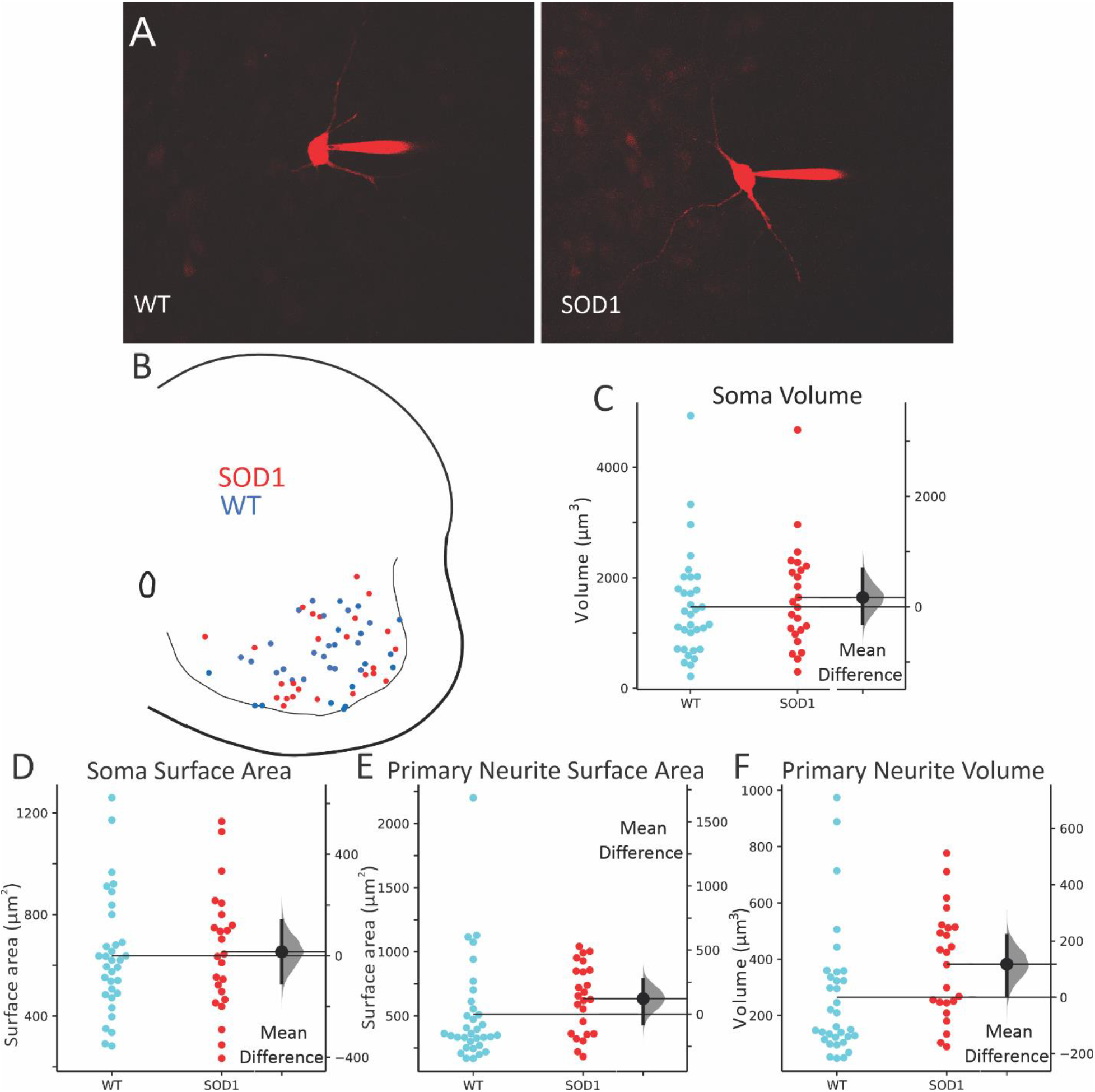
GlyT2 interneurons from WT and SOD1 mice were patched throughout the ventral horn. Images of typical interneurons, filled with Texas Red as they were recorded in **A**, and locations of all patched interneurons in **B** (WT = blue circles, SOD1 = red circles). Consistent with previous results, soma volume (**C**) and soma surface area (**D**) were unchanged in patched neurons, while total surface area and total volume of primary neurites (**E** and **F**) were modestly larger in SOD1 interneurons than in WT. Effect sizes are shown to the right of each swarm plot, with distribution of the 95% CI of the effect sizes in grey to the right of the mean difference (SOD1 – WT).

### Electrophysiology of glycinergic interneurons

Whole cell patch clamp revealed very different intrinsic properties of ventral glycinergic interneurons. Recording was performed on 59 ventral interneurons from throughout the ventral horn in transverse lumbar spinal cord slices from P6-10 mice. SOD1 inhibitory interneurons (n = 25 from 10 mice) were found to have diminished intrinsic excitability compared to WT interneurons (n = 34 from 15 mice). Measurements of intrinsic excitability included depolarized onset and peak voltage of PICs, resulting in a small change in threshold for action potential firing, and the current at firing onset (I-ON) and offset (I-OFF) as shown in **Figure 3**. In voltage clamp, slowly depolarizing voltage ramps were used to measure PICs (typical leak subtracted traces shown in Fig 3A). Voltage sensitivity is determined by measuring voltage at PIC onset and peak. Both PIC onset and peak were significantly depolarized in SOD1 interneurons, while amplitude of the PIC was unchanged. Depolarizing current ramps were used to measure the input-output relationship of the neurons in current clamp. Inhibitory interneurons from SOD1 mice were found to have significantly higher threshold voltage than WT interneurons (Fig 3E-F). This shift is likely driven by changes in PICs. The current at firing onset, or I-ON, and firing offset (I-OFF) was similarly larger in SOD1 interneurons. The AHP appeared to be shorter duration in SOD1 interneurons, however corresponding changes in the decay rate (tau) were not present. The total AHP duration and the duration at half amplitude were slightly shorter in SOD1 neurons, as shown in Fig 3J and **Table 2**, though amplitude of the AHP was unchanged. While changes in the PIC and threshold indicate less excitability in SOD1 interneurons, the shortened AHP suggests an increased ability to fire at higher rates in SOD1 interneurons, a property that is associated with increased excitability. However, no changes in firing rates were detected in SOD1 interneurons. All other properties were found to be similar in SOD1 and WT glycinergic interneurons, including maximum firing rates, sag/rebound currents (I_H_), action potential parameters and membrane properties (see Table 2 for all electrical properties). See discussion for further interpretation of these findings.

**Figure 3:**
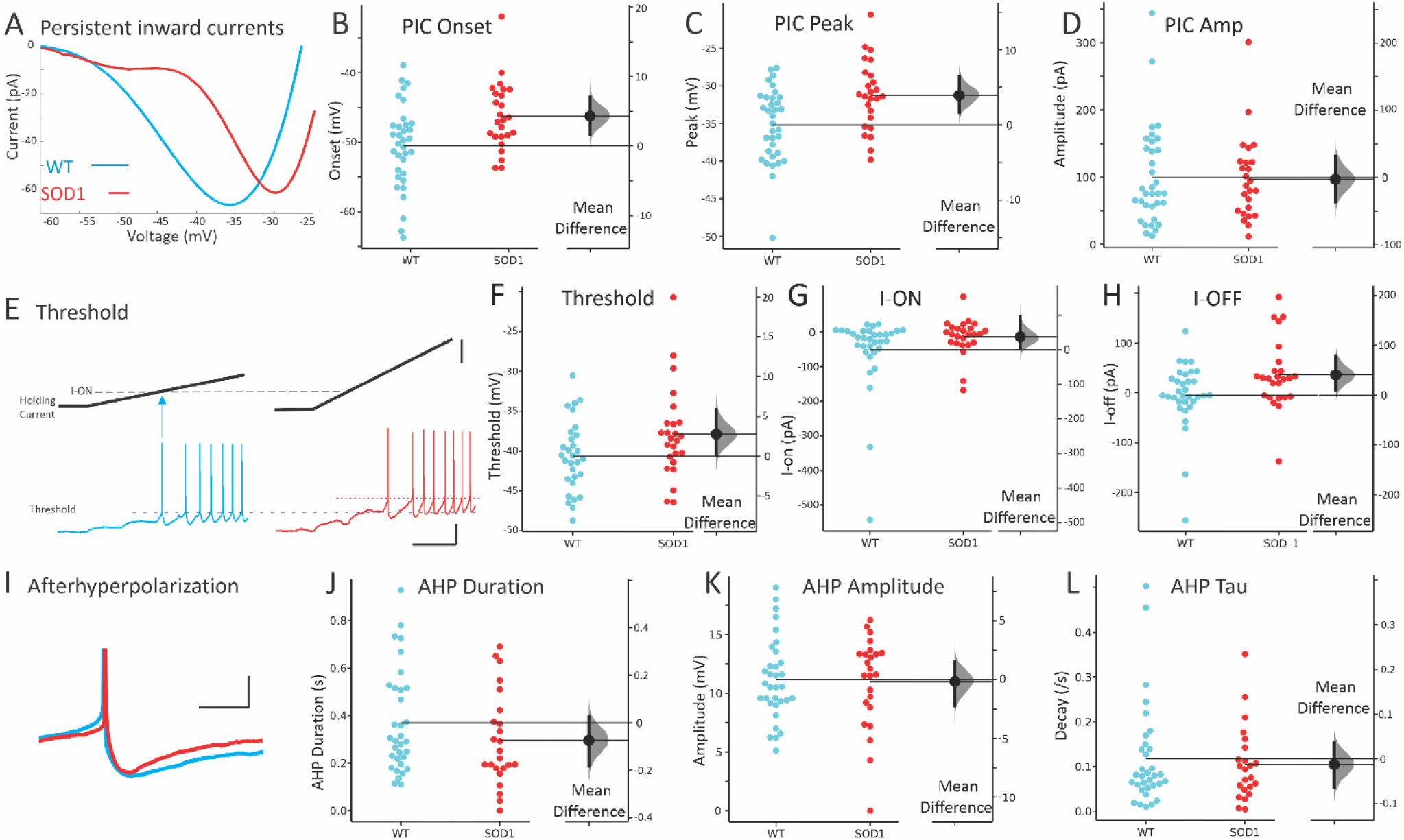
Electrophysiology of SOD1 glycinergic interneurons. The most prominent difference in SOD1 interneurons was the shift in voltage dependence of PICs. (**A**) Representative, leak-subtracted current-voltage relationship of PICs from WT (P8) and SOD1 (P6) interneurons. (**B-D**) Mean onset, peak and amplitude of PICs in WT (blue symbols on left) and SOD1 interneurons (red symbols on right). (**E-F**) Threshold for action potentials evoked with current ramps was higher in SOD1 interneurons (compare dotted lines indicating voltage threshold). Starting potential for both interneurons was −65 mV. The current at firing onset and offset (I-ON and I-OFF, respectively) was also increased as shown in **G-H**. (**I**) The duration of the AHP in SOD1 interneurons appeared shorter, as shown in a representative P8 WT and P7 SOD1 interneuron and cumulative data for total duration of AHP in **J**. However, neither the AHP amplitude nor the AHP decay time constant, tau, were significantly altered in SOD1 glycinergic interneurons, as shown in **K-L**. Vertical scale bars in E: top = 50 pA, and bottom = 20 mV. Horizontal scale bar in **E** = 0.5s and all scale bars apply to both left and right panels. Vertical scale bar in H = 10 mV (APs were truncated from image), and horizontal scale bar = 50 ms.

**Table 2:**
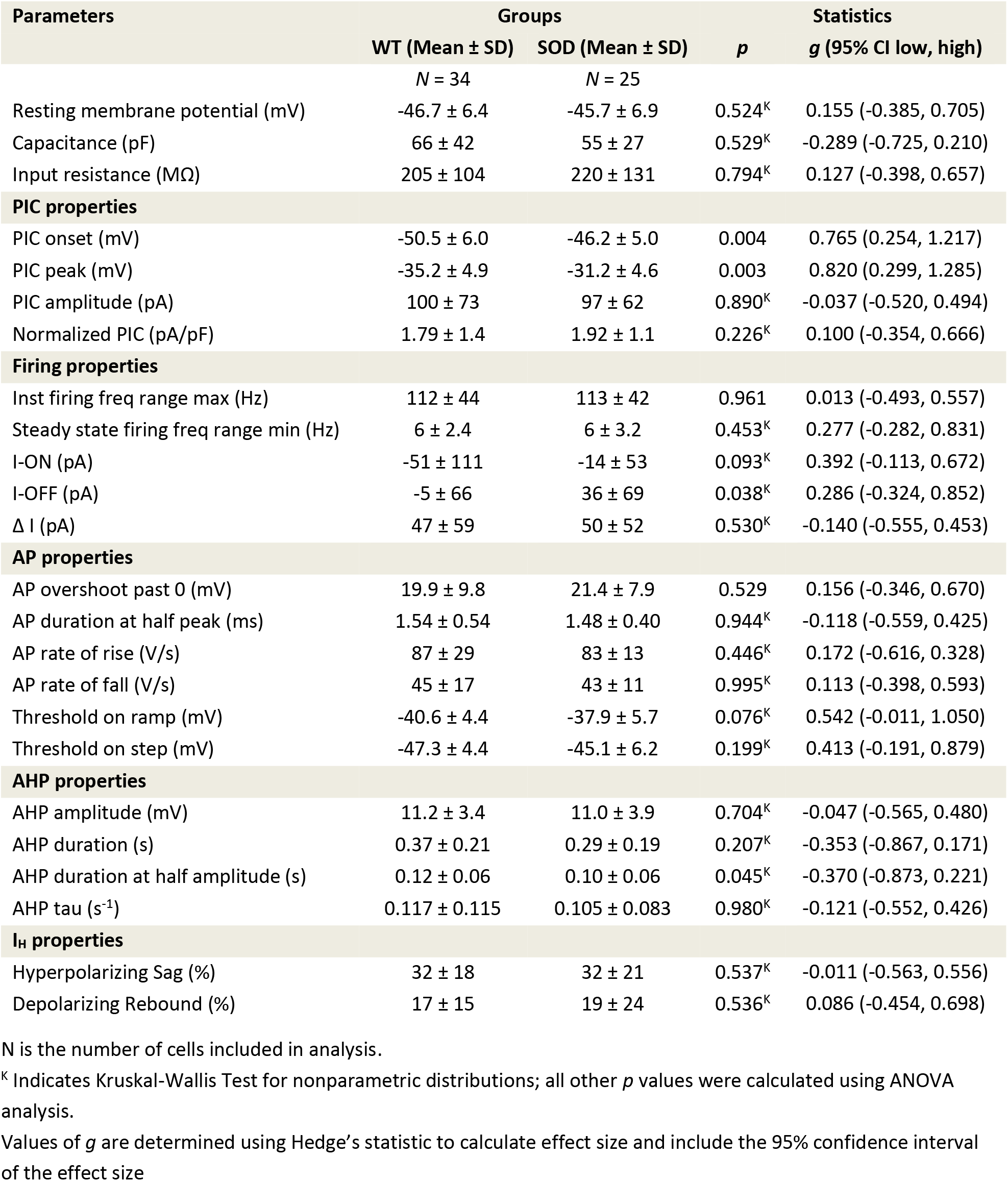
Electrophysiological properties of GlyT2-GFP+ interneurons in WT and SOD1 animals.

### Regional analysis of glycinergic interneurons

Since ventral glycinergic interneurons are not a homogenous population, we tested for regional differences. Interneurons were divided into three groups based on location: 1) the “ventral RC region”: within 100 μm of the ventral white matter where Renshaw cells are located, 2) lamina IX, and 3) the “intermediate region”: ventral horn region including lamina VII and VIII, excluding the previous two regions. These regions are represented in **Figure 4**. The first region corresponds to the location of RCs (but could contain other types of glycinergic interneurons). The second region contains interneurons that are interspersed with the MNs within the motor pools, likely to be composed of premotor interneurons (more on this in the discussion). The third region contains interneurons that are a mix of premotor interneurons and other classes of ventral interneurons. There was overlap of the first two regions, so interneurons within the ventral most 100 μm that also were located within motor pools were included in both the ventral RC region and lamina IX, as shown in open circles in Figure 4.

**Figure 4:**
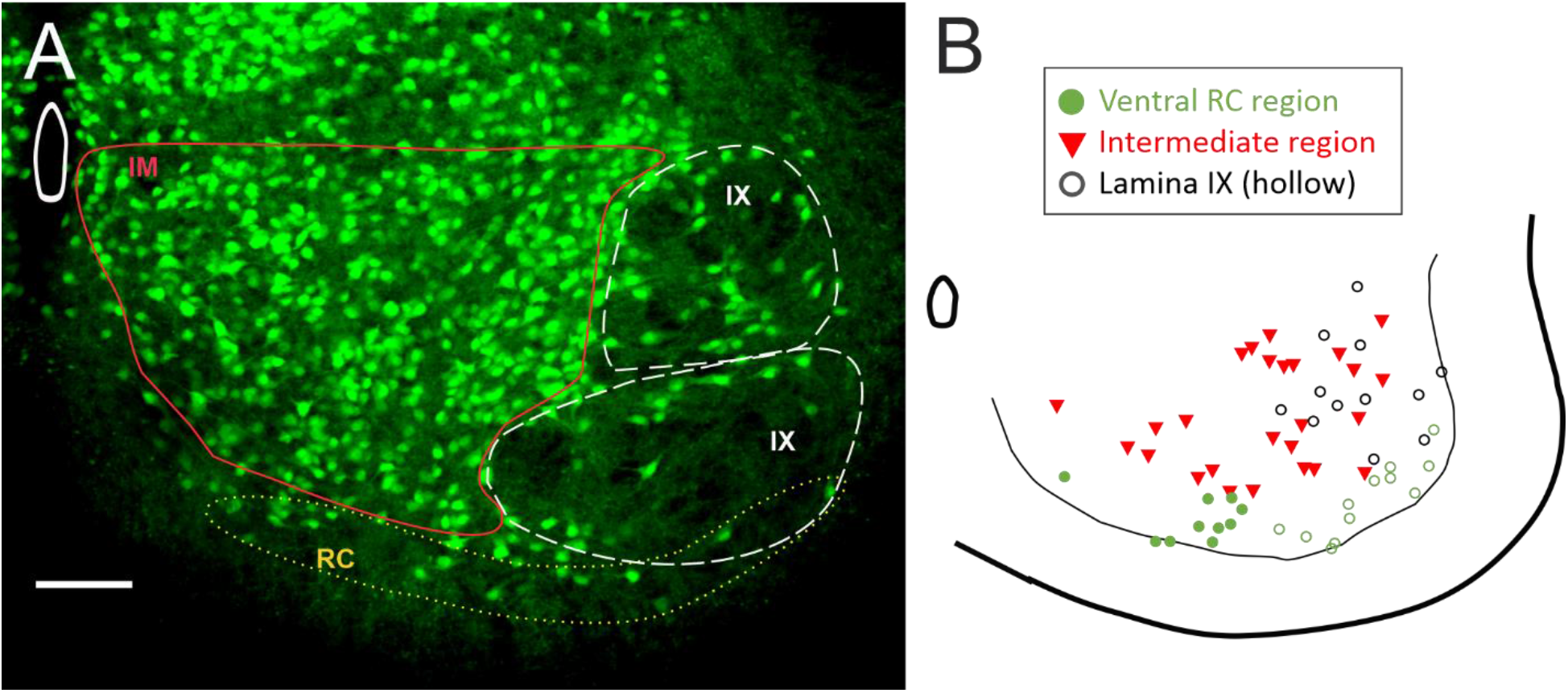
Regional divisions of interneurons. (**A**) Three regions were used for analysis of regional differences in glycinergic interneurons. (**B**) Map of the patched interneurons coded for regions. Ventral region with RC cells (green circles) overlapped with lamina IX (hollow circles). The remaining interneurons that were not within the RC region or lamina IX were grouped into an intermediate region of lamina VII and VIII (red triangles). Scale bar in **A** = 100 μm.

#### GlyT2 interneurons in Ventral RC region

Within the ventral region where RCs are located, we found that glycinergic SOD1 interneurons there showed the most significant changes in both electrophysiology and morphology. In patch clamped interneurons from this region (N = 8 WT from 7 mice and 13 SOD1 from 9 mice), there was a significant depolarization in PIC onset and peak in glycinergic SOD1 interneurons compared to WT interneurons in the same location. Additionally, primary neurites were found to be longer and larger in volume and surface area in SOD1 interneurons in the ventral-most 100 μm. Unpatched GlyT2+ SOD1 interneurons in this region (188 WT interneurons from 6 WT mice and 225 SOD1 interneurons from 6 SOD1 mice), analyzed from fixed spinal cords were smaller than WT glycinergic interneurons. These changes (summarized in **Figure 5** and **Table 3**) show that glycinergic neurons in the region where RCs are located are very affected by the SOD1 mutation. Primary neurites, soma volume, PIC onset and PIC peak were all as affected or more affected in the RC region than the larger group of ventral horn interneurons (for example compare primary neurite measurements in Fig 2E-F and Fig 5G-H).

**Figure 5:**
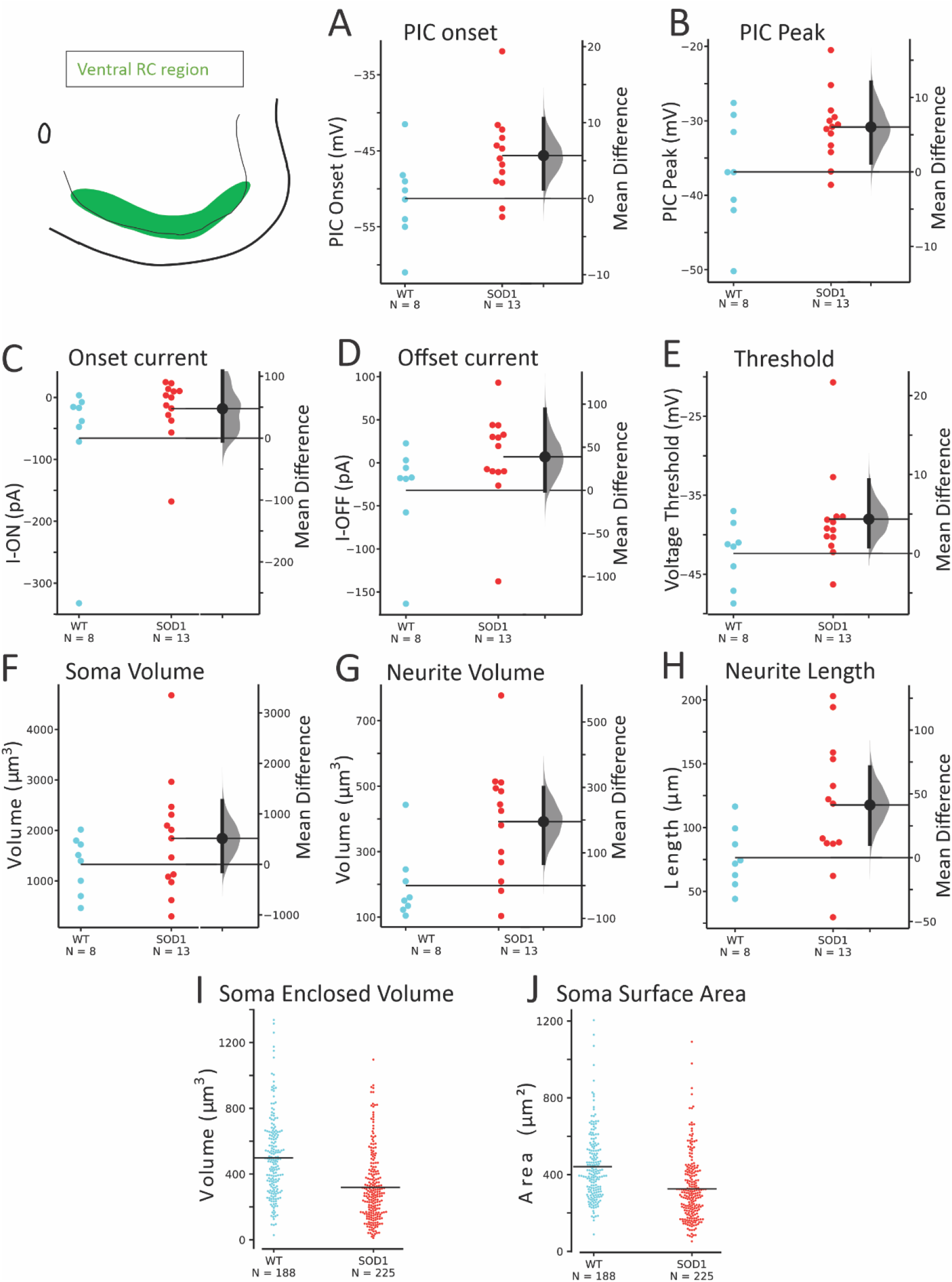
SOD1 GlyT2 interneurons in the ventral RC region were altered in electrophysiology and morphology. Both onset (**A**) and peak (**B**) of the PIC were depolarized. Threshold voltage and current at firing onset and offset were more variable as shown by the effect sizes (**C-E**). Soma volume of patched neurons was not significantly altered but neurites were larger in total volume and total length (**F-H**) in SOD1 interneurons. Larger numbers of unpatched neurons were analyzed for soma volume (**I**) and surface area (**J**) in the RC region. A linear mixed model (including inter-mouse variability) revealed that soma volume was significantly smaller in SOD1 mice in this region.

**Table 3:**
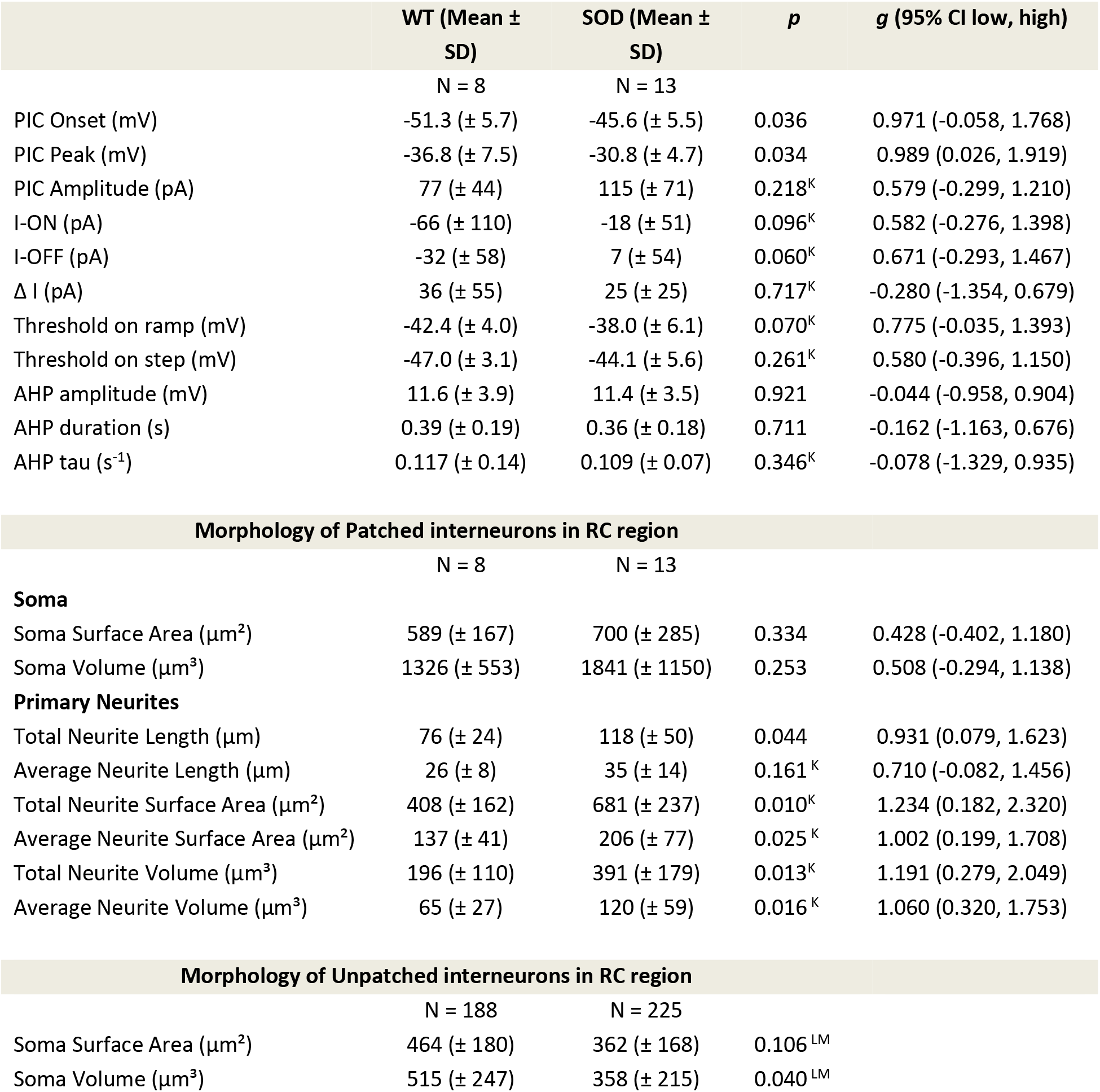
Electrophysiological properties of WT and SOD1 GlyT2-GFP+ interneurons in ventral RC region. ^K^ Indicates Kruskal-Wallis Test for nonparametric distributions; ^LM^ Indicates linear mixed effects model; all others were ANOVA analysis. Values of *g* are determined using Hedge’s statistic to calculate effect size and include the 95% confidence interval of the effect size

| | WT (Mean ± SD) | SOD (Mean ± SD) | *p* | *g* (95% CI low, high) |
| --- | --- | --- | --- | --- |
|  | N = 8 | N = 13 |  |  |
| PIC Onset (mV) | -51.3 (± 5.7) | -45.6 (± 5.5) | 0.036 | 0.971 (-0.058, 1.768) |
| PIC Peak (mV) | -36.8 (± 7.5) | -30.8 (± 4.7) | 0.034 | 0.989 (0.026, 1.919) |
| PIC Amplitude (pA) | 77 (± 44) | 115 (± 71) | 0.218<sup>K</sup> | 0.579 (-0.299, 1.210) |
| I-ON (pA) | -66 (± 110) | -18 (± 51) | 0.096<sup>K</sup> | 0.582 (-0.276, 1.398) |
| I-OFF (pA) | -32 (± 58) | 7 (± 54) | 0.060<sup>K</sup> | 0.671 (-0.293, 1.467) |
| $\Delta I$ (pA) | 36 (± 55) | 25 (± 25) | 0.717<sup>K</sup> | -0.280 (-1.354, 0.679) |
| Threshold on ramp (mV) | -42.4 (± 4.0) | -38.0 (± 6.1) | 0.070<sup>K</sup> | 0.775 (-0.035, 1.393) |
| Threshold on step (mV) | -47.0 (± 3.1) | -44.1 (± 5.6) | 0.261<sup>K</sup> | 0.580 (-0.396, 1.150) |
| AHP amplitude (mV) | 11.6 (± 3.9) | 11.4 (± 3.5) | 0.921 | -0.044 (-0.958, 0.904) |
| AHP duration (s) | 0.39 (± 0.19) | 0.36 (± 0.18) | 0.711 | -0.162 (-1.163, 0.676) |
| AHP tau (s<sup>-1</sup>) | 0.117 (± 0.14) | 0.109 (± 0.07) | 0.346<sup>K</sup> | -0.078 (-1.329, 0.935) |
| Morphology of Patched interneurons in RC region |  |  |  |  |
|  | N = 8 | N = 13 |  |  |
| **Soma** |  |  |  |  |
| Soma Surface Area ($\mu\text{m}^2$) | 589 (± 167) | 700 (± 285) | 0.334 | 0.428 (-0.402, 1.180) |
| Soma Volume ($\mu\text{m}^3$) | 1326 (± 553) | 1841 (± 1150) | 0.253 | 0.508 (-0.294, 1.138) |
| **Primary Neurites** |  |  |  |  |
| Total Neurite Length ($\mu\text{m}$) | 76 (± 24) | 118 (± 50) | 0.044 | 0.931 (0.079, 1.623) |
| Average Neurite Length ($\mu\text{m}$) | 26 (± 8) | 35 (± 14) | 0.161<sup>K</sup> | 0.710 (-0.082, 1.456) |
| Total Neurite Surface Area ($\mu\text{m}^2$) | 408 (± 162) | 681 (± 237) | 0.010<sup>K</sup> | 1.234 (0.182, 2.320) |
| Average Neurite Surface Area ($\mu\text{m}^2$) | 137 (± 41) | 206 (± 77) | 0.025<sup>K</sup> | 1.002 (0.199, 1.708) |
| Total Neurite Volume ($\mu\text{m}^3$) | 196 (± 110) | 391 (± 179) | 0.013<sup>K</sup> | 1.191 (0.279, 2.049) |
| Average Neurite Volume ($\mu\text{m}^3$) | 65 (± 27) | 120 (± 59) | 0.016<sup>K</sup> | 1.060 (0.320, 1.753) |

**Table 3:** Electrophysiological properties of WT and SOD1 GlyT2-GFP+ interneurons in ventral RC region.
|  |  |  |  |  |
| --- | --- | --- | --- | --- |
|  | N = 188 | N = 225 |  |  |
| Soma Surface Area ($\mu\text{m}^2$) | 464 (± 180) | 362 (± 168) | 0.106<sup>LM</sup> | |
| Soma Volume ($\mu\text{m}^3$) | 515 (± 247) | 358 (± 215) | 0.040<sup>LM</sup> | |

#### GlyT2 interneurons in Lamina IX

The SOD1 interneurons located in lamina IX, intermixed with MNs were modestly altered in excitability and morphology. As shown in **Figure 6** and **Table 4**, PIC onset was depolarized in SOD1 interneurons in this region, compared to interneurons in this region from WT mice (n = 14 WT interneurons from 8 mice and 11 SOD1 interneurons from 9 mice). Longer total length of primary neurites was also found in patched interneurons in this region. Within the larger number of unpatched interneurons (310 WT interneurons from 6 WT mice and 258 SOD1 interneurons from 6 SOD1 mice), there was no significant change in soma size.

**Figure 6:**
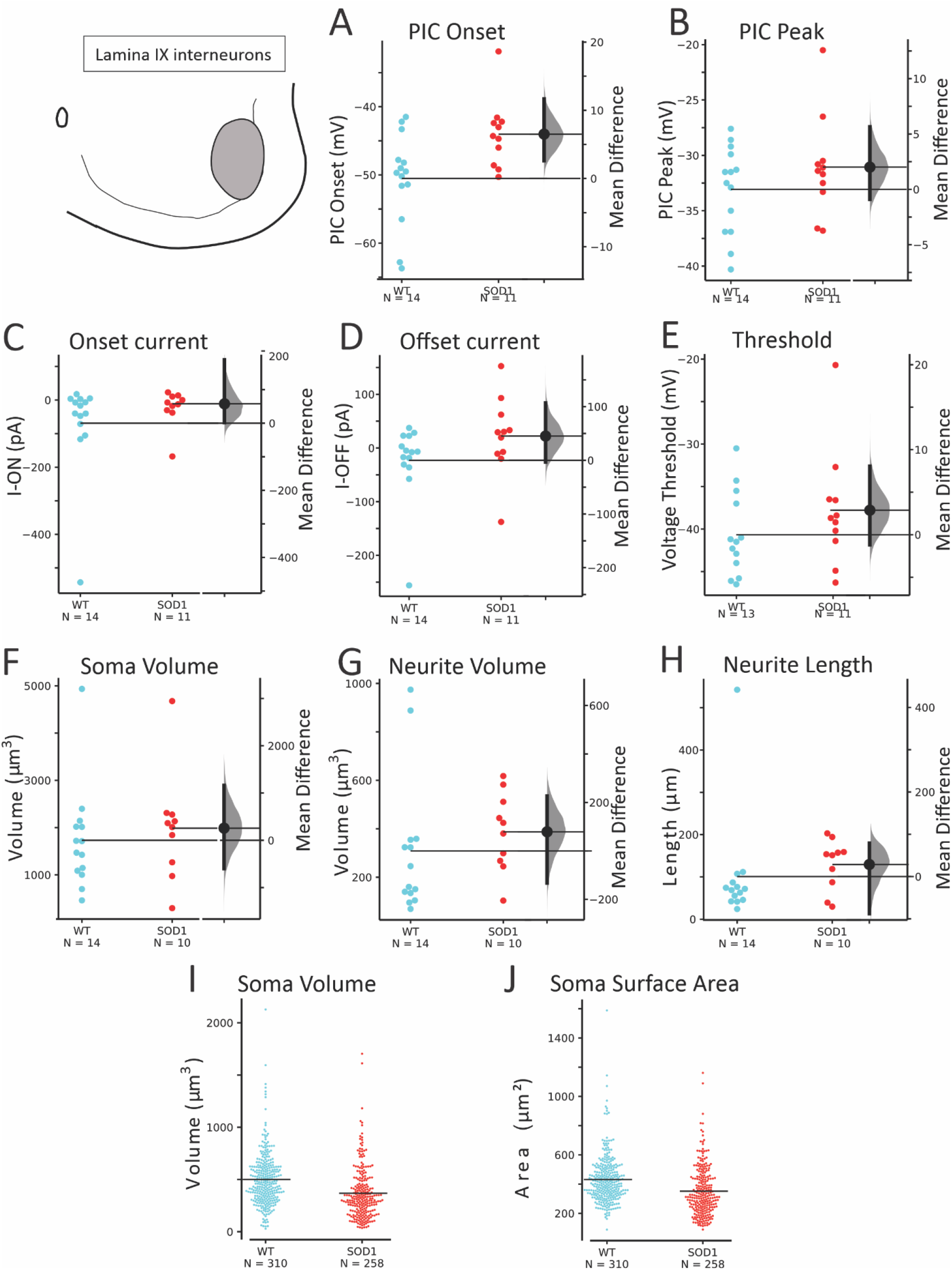
Modest changes in electrophysiology were present in glycinergic SOD1 interneurons in lamina IX. PIC onset (**A**) was depolarized in SOD1 interneurons without reliable changes in PIC peak (**B**), voltage threshold, I-ON or I-OFF (**C-E**). Soma (**F**) and primary neurites (**G-H**) did not show consistent changes. Unpatched glycinergic neurons were not significantly altered in soma enclosed volume (**I**) and surface area (**J**) in SOD1 interneurons compared to WT in Lamina IX.

**Table 4:** Electrophysiological properties of WT and SOD1 GlyT2-GFP+ interneurons in lamina IX.

| | WT (Mean ± SD) | SOD (Mean ± SD) | *p* | *g* (95% CI low, high) |
| --- | --- | --- | --- | --- |
|  | N = 14 | N = 11 |  |  |
| PIC Onset (mV) | -50.5 (± 6.7) | -44.0 (± 5.0) | 0.013 | 1.046 (0.340, 1.652) |
| PIC Peak (mV) | -33.1 (± 3.9) | -31.1 (± 4.5) | 0.248 | 0.462 (-0.349, 1.165) |
| PIC Amplitude (pA) | 116 (± 81) | 98 (± 76) | 0.412<sup>K</sup> | -0.218 (-0.944, 0.655) |
| I-ON (pA) | -69 (± 143) | -11 (± 64) | 0.155<sup>K</sup> | 0.485 (-0.439, 0.928) |
| I-OFF (pA) | -23 (± 72) | 22 (± 73) | 0.080<sup>K</sup> | 0.602 (-0.401, 1.207) |
| $\Delta$ I (pA) | 48 (± 76) | 33 (± 30) | 0.702<sup>K</sup> | -0.226 (-0.564, 1.021) |
| Threshold on ramp (mV) | -40.7 (± 4.9) | -37.8 (± 6.8) | 0.244 | 0.473 (-0.370, 1.209) |
| Threshold on step (mV) | -47.4 (± 4.2) | -43.9 (± 7.4) | 0.112<sup>K</sup> | 0.595 (-0.437, 1.377) |
| AHP amplitude (mV) | 11.4 (± 3.9) | 10.2 (± 3.5) | 0.421 | -0.319 (-1.09, 0.514) |
| AHP duration (s) | 0.41 (± 0.24) | 0.25 (± 0.16) | 0.062 | -0.764 (-1.509, 0.053) |
| AHP tau (s<sup>-1</sup>) | 0.129 (± 0.118) | 0.076 (± 0.067) | 0.274<sup>K</sup> | -0.511 (-1.088, 0.390) |

| Morphology of Patched interneurons in lamina IX |  |  |  |  |
| --- | --- | --- | --- | --- |
|  | N = 14 | N = 10 |  |  |
| **Soma** |  |  |  |  |
| Soma Surface Area ($\mu\text{m}^2$) | 703 (± 242) | 734 (± 284) | 0.771 | 0.118 (-0.740, 0.959) |
| Soma Volume ($\mu\text{m}^3$) | 1732 (± 1080) | 1989 (± 1149) | 0.412<sup>K</sup> | 0.224 (-0.620, 1.076) |
| **Primary Neurites** |  |  |  |  |
| Total Neurite Length ($\mu\text{m}$) | 101 (± 130) | 129 (± 60) | 0.046<sup>K</sup> | 0.258 (-0.610, 1.855) |
| Average Neurite Length ($\mu\text{m}$) | 29 (± 24) | 37 (± 16) | 0.069<sup>K</sup> | 0.335 (-0.544, 1.442) |
| Total Neurite Surface Area ($\mu\text{m}^2$) | 545 (± 524) | 721 (± 293) | 0.089<sup>K</sup> | 0.384 (-0.522, 1.722) |
| Average Neurite Surface Area ($\mu\text{m}^2$) | 164 (± 99) | 208 (± 80) | 0.128<sup>K</sup> | 0.466 (-0.435, 1.381) |
| Total Neurite Volume ($\mu\text{m}^3$) | 309 (± 283) | 388 (± 161) | 0.114<sup>K</sup> | 0.316 (-0.498, 1.474) |
| Average Neurite Volume ($\mu\text{m}^3$) | 95 (± 65) | 113 (± 46) | 0.178<sup>K</sup> | 0.307 (-0.496, 1.233) |

| Morphology of Unpatched interneurons in lamina IX |  |  |  |  |
| --- | --- | --- | --- | --- |
|  | N = 310 | N = 258 |  |  |
| Soma Surface Area ($\mu\text{m}^2$) | 459 (± 164) | 372 (± 169) | 0.069<sup>LM</sup> | |
| Soma Volume ($\mu\text{m}^3$) | 525 (± 257) | 392 (± 248) | 0.067<sup>LM</sup> | |

<sup>K</sup> Indicates Kruskal-Wallis Test for nonparametric distributions; <sup>LM</sup> Indicates linear mixed effects model; all others were ANOVA analysis. Values of *g* are determined using Hedge's statistic to calculate effect size and include the 95% confidence interval of the effect size

#### GlyT2 interneurons in the Intermediate region

The third regional group of glycinergic interneurons was defined as the intermediate region. These interneurons were located in laminae VII and VIII, excluding the ventral 100 μm and lamina IX. Within this area, 16 WT and 8 SOD1 interneurons were patched from 11 WT and 5 SOD1 mice respectively, and 655 unpatched interneurons were analyzed from 6 WT mice and 812 unpatched SOD1 interneurons from 6 SOD1 mice. As shown in **Figure 7** and **Table 5**, the PIC peak was the only parameter that was altered in SOD1 neurons in this region. Overall, the GlyT2 interneurons located here were the least affected by the SOD1 mutation.

**Figure 7:**
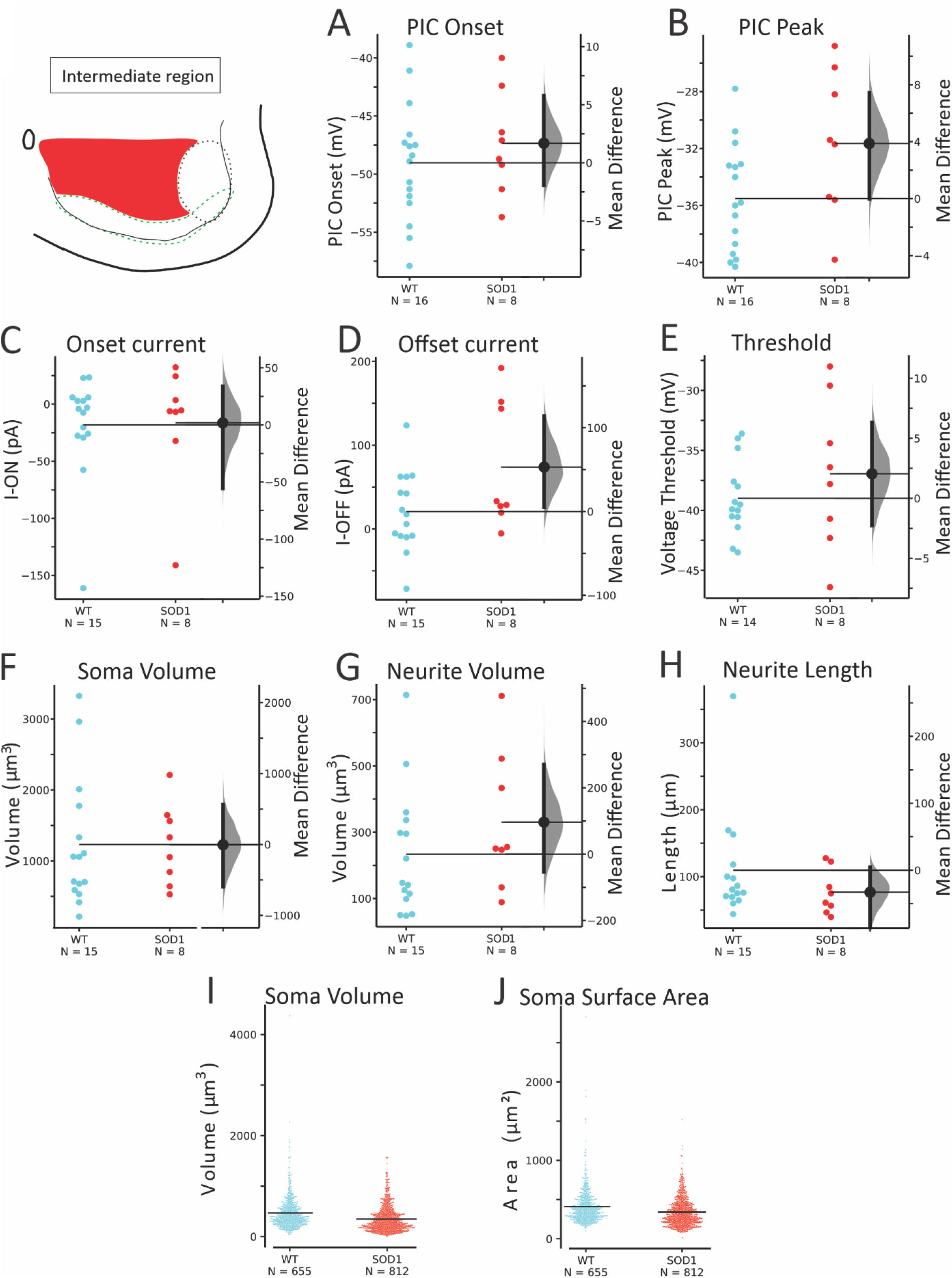
SOD1 interneurons in the intermediate regions of laminae VII and VIII were largely unaffected. PIC onset (**A**) was not significantly different, though PIC peak (**B**) was marginally more depolarized in SOD1 neurons. Voltage and current threshold (**C-E**) were not significantly affected. Morphology of soma (**F**) and neurites (**G-H**) in patched SOD1 interneurons were unchanged, and larger numbers of unpatched neurons in this region (**I-J**) were not significantly smaller.

**Table 5:**
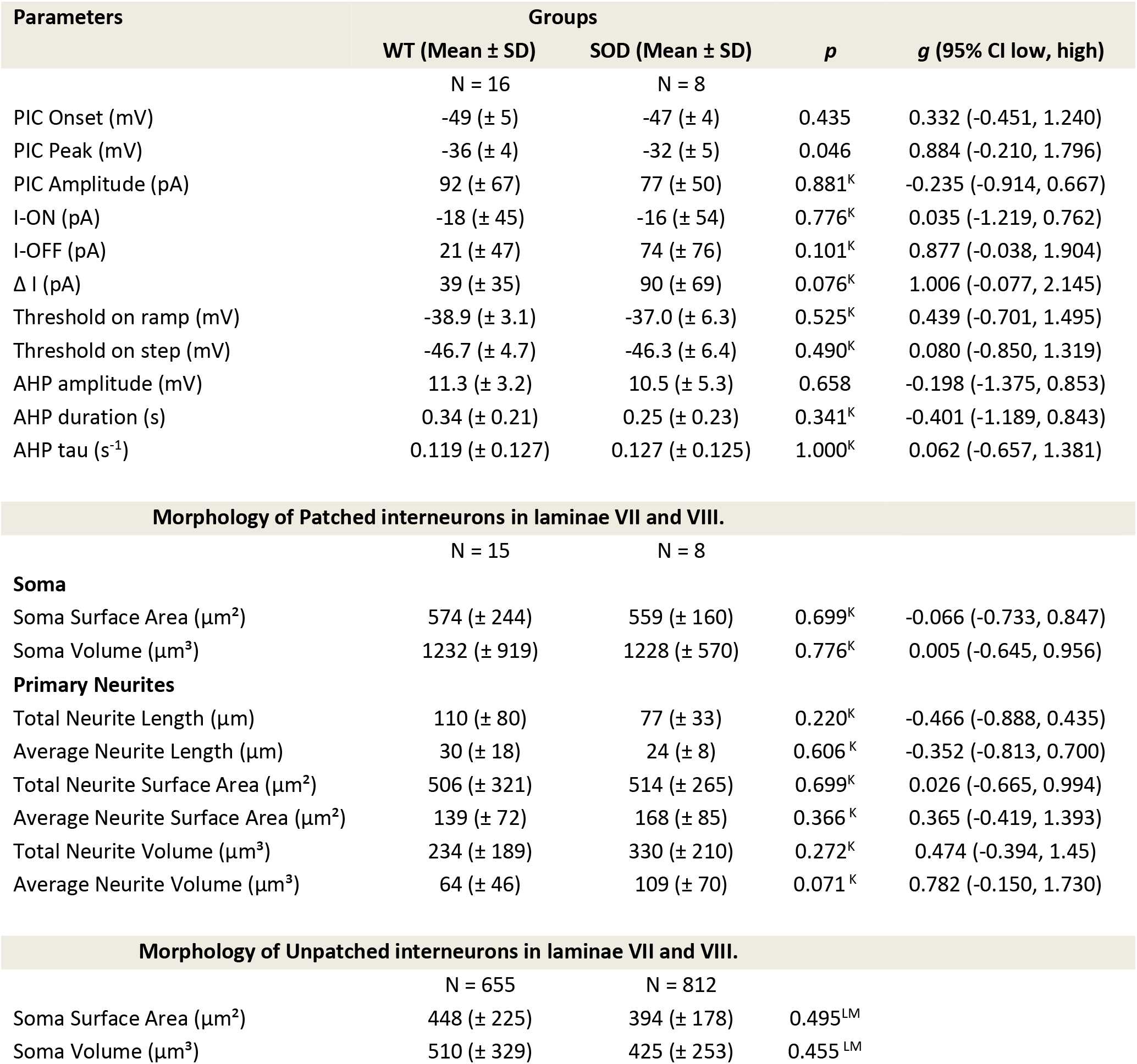
Electrophysiological properties of WT and SOD1 GlyT2-GFP+ interneurons in Intermediate Region of lamina VII and VIII. ^K^ Indicates Kruskal-Wallis Test for nonparametric distributions; ^LM^ Indicates linear mixed effects model; all others were ANOVA analysis. Values of *g* are determined using Hedge’s statistic to calculate effect size and include the 95% confidence interval of the effect size

| Parameters | Groups |  | *p* | *g* (95% CI low, high) |
| --- | --- | --- | --- | --- |
|  | WT (Mean ± SD) | SOD (Mean ± SD) |  |  |
|  | N = 16 | N = 8 |  |  |
| PIC Onset (mV) | -49 (± 5) | -47 (± 4) | 0.435 | 0.332 (-0.451, 1.240) |
| PIC Peak (mV) | -36 (± 4) | -32 (± 5) | 0.046 | 0.884 (-0.210, 1.796) |
| PIC Amplitude (pA) | 92 (± 67) | 77 (± 50) | 0.881<sup>K</sup> | -0.235 (-0.914, 0.667) |
| I-ON (pA) | -18 (± 45) | -16 (± 54) | 0.776<sup>K</sup> | 0.035 (-1.219, 0.762) |
| I-OFF (pA) | 21 (± 47) | 74 (± 76) | 0.101<sup>K</sup> | 0.877 (-0.038, 1.904) |
| Δ I (pA) | 39 (± 35) | 90 (± 69) | 0.076<sup>K</sup> | 1.006 (-0.077, 2.145) |
| Threshold on ramp (mV) | -38.9 (± 3.1) | -37.0 (± 6.3) | 0.525<sup>K</sup> | 0.439 (-0.701, 1.495) |
| Threshold on step (mV) | -46.7 (± 4.7) | -46.3 (± 6.4) | 0.490<sup>K</sup> | 0.080 (-0.850, 1.319) |
| AHP amplitude (mV) | 11.3 (± 3.2) | 10.5 (± 5.3) | 0.658 | -0.198 (-1.375, 0.853) |
| AHP duration (s) | 0.34 (± 0.21) | 0.25 (± 0.23) | 0.341<sup>K</sup> | -0.401 (-1.189, 0.843) |
| AHP tau (s<sup>-1</sup>) | 0.119 (± 0.127) | 0.127 (± 0.125) | 1.000<sup>K</sup> | 0.062 (-0.657, 1.381) |
| Morphology of Patched interneurons in laminae VII and VIII. |  |  |  |  |
|  | N = 15 | N = 8 |  |  |
| **Soma** |  |  |  |  |
| Soma Surface Area (μm<sup>2</sup>) | 574 (± 244) | 559 (± 160) | 0.699<sup>K</sup> | -0.066 (-0.733, 0.847) |
| Soma Volume (μm<sup>3</sup>) | 1232 (± 919) | 1228 (± 570) | 0.776<sup>K</sup> | 0.005 (-0.645, 0.956) |
| **Primary Neurites** |  |  |  |  |
| Total Neurite Length (μm) | 110 (± 80) | 77 (± 33) | 0.220<sup>K</sup> | -0.466 (-0.888, 0.435) |
| Average Neurite Length (μm) | 30 (± 18) | 24 (± 8) | 0.606<sup>K</sup> | -0.352 (-0.813, 0.700) |
| Total Neurite Surface Area (μm<sup>2</sup>) | 506 (± 321) | 514 (± 265) | 0.699<sup>K</sup> | 0.026 (-0.665, 0.994) |
| Average Neurite Surface Area (μm<sup>2</sup>) | 139 (± 72) | 168 (± 85) | 0.366<sup>K</sup> | 0.365 (-0.419, 1.393) |
| Total Neurite Volume (μm<sup>3</sup>) | 234 (± 189) | 330 (± 210) | 0.272<sup>K</sup> | 0.474 (-0.394, 1.45) |
| Average Neurite Volume (μm<sup>3</sup>) | 64 (± 46) | 109 (± 70) | 0.071<sup>K</sup> | 0.782 (-0.150, 1.730) |

**Table 5:** Electrophysiological properties of WT and SOD1 GlyT2-GFP+ interneurons in Intermediate Region of lamina VII and VIII.
|  |  |  |  |
| --- | --- | --- | --- |
|  | N = 655 | N = 812 |  |
| Soma Surface Area (μm<sup>2</sup>) | 448 (± 225) | 394 (± 178) | 0.495<sup>LM</sup> |
| Soma Volume (μm<sup>3</sup>) | 510 (± 329) | 425 (± 253) | 0.455<sup>LM</sup> |

Taken together, these results show that glycinergic interneurons are affected by the SOD1 mutation, particularly those in the ventral-most region of the spinal cord. These neurons are altered very early in life in both electrophysiology and morphology.

## Discussion

This study shows for the first time that glycinergic interneurons are depressed in excitability in the SOD1 mouse model and may be contributing to ALS pathology. Alterations in both morphology and excitability of glycinergic interneurons occur at a very early, presymptomatic stage, and are particularly prominent in those interneurons located within 100 μm of the ventral white matter. We speculate that these shifts in interneuron excitability and morphology could have functional consequences that alter inhibition of MNs, modify motor output, and could even provide an early biomarker of ALS.

### Intrinsic and synaptic hyperexcitability

A core issue this study brings to light is potential long-term disruption to the balance of excitatory and inhibitory synaptic transmission in ALS. Spinal cord and cortex both show an imbalance between excitatory and inhibitory synaptic transmission in SOD^G93A^ mice (Avossa et al., 2006; Saba et al., 2016). Clinically, ALS patients show evidence of altered synaptic activity as well, including a reduction in inhibition. A reduction of glycinergic receptor binding in the anterior gray matter of the spinal cord (Hayashi et al., 1981; Whitehouse et al., 1983) has been demonstrated along with abnormal glycine and gamma-aminobutyric acid (GABA) levels in blood serum (Malessa et al., 1991; Niebroj-Dobosz and Janik, 1999). Electrophysiology also suggests disruption in spinal inhibitory circuits in ALS patients (Raynor and Shefner, 1994; Shefner and Logigian, 1998; Sangari et al., 2016; Özyurt et al., 2020). An interesting question is whether the neurons are altered in excitability only within the motor circuit (i.e., corticospinal neurons and the synaptically-connected spinal inhibitory and cholinergic interneurons, and MNs), or if disturbances are more widespread throughout the nervous system. In the spinal cord, interneurons that serve as conduits between vulnerable corticospinal and spinal MNs include cholinergic interneurons, Ia inhibitory interneurons and RCs. There is abundant evidence of disruption in these cholinergic, GABAergic and glycinergic synapses from animal models (Martin et al., 2007; Chang and Martin, 2009b; Pullen and Athanasiou, 2009; Hossaini et al., 2011; Herron and Miles, 2012; Casas et al., 2013; Saxena et al., 2013; Wootz et al., 2013; Milan et al., 2015; Dukkipati et al., 2016; Medelin et al., 2016). Similarly, in the cortex, neurons that are not themselves vulnerable to neurodegeneration in ALS also exhibit altered excitability and morphology in a parallel time course to the vulnerable corticospinal neurons (Clark et al., 2017, 2018; Kim et al., 2017). The changes in excitability of other neuronal populations must contribute to the imbalance of both excitatory and inhibitory neurotransmission that has been clearly demonstrated in ALS patients and animal models.

A case for hyperexcitability and glutamate excitotoxicity in ALS pathogenesis has been more controversial. Altered intrinsic excitability has been consistently reported in both corticospinal projection neurons and spinal MNs, but these perturbations appear to fluctuate based on age/disease progression. Corticospinal projection neurons have increased firing and other signs of hyperexcitability during the first week of postnatal development (Pieri et al., 2009; Saba et al., 2016, 2019; Kim et al., 2017), but excitability is normal in young and adult pre-symptomatic ages (Kim et al., 2017; Saba et al., 2019), and may become hypoexcitable at the age of symptom onset (Saba et al., 2019). Similarly, spinal MNs show intrinsic hyperexcitability at very early (embryonic) stages (Kuo et al., 2004, 2005; Martin et al., 2013). At postnatal ages (similar to the inhibitory interneurons in this study), spinal MNs do *not* have an altered threshold, rheobase or frequency-current relationship, though other abnormalities are present in PICs, action potential duration, AHP and dendritic Ca^2+^ entry (Pambo-Pambo et al., 2009; Quinlan et al., 2011, 2015; Leroy et al., 2014). Thus, at the age studied here, inhibitory interneurons are affected differently than the vulnerable spinal MNs. At symptom onset, MNs may finally succumb to hypoexcitability, though not all studies agree on this point (Delestrée et al., 2014; Martinez-Silva et al., 2018; Jensen et al., 2020). Notably, MNs derived from ALS patients’ induced pluripotent stem cells also show initial hyperexcitability followed by hypoexcitability (Wainger et al., 2014; Devlin et al., 2015), and increasing excitability of MNs enhanced neuroprotection rather than neurodegeneration (Saxena et al., 2013). A unifying theme throughout the findings is that vulnerable MNs show excessive homeostatic gain in response to perturbation (Kuo et al., 2020), and not only excitability but other cellular properties fluctuate over the lifespan of the animal (Irvin et al., 2015). In light of that, whether inhibitory interneurons are less excitable at other ages should be explored in future studies to fully characterize their contribution to neurodegenerative processes.

### Changes in neuron morphology

In ALS models, altered morphology in spinal MNs occurs so early that it could be viewed as a deficit in normal development. Embryonically, SOD1 MNs have shorter projections but this is reversed during postnatal development. At 1-2 weeks of age, SOD1 MNs begin to show more dendritic branching and larger soma sizes than WT MNs (Amendola and Durand, 2008; Filipchuk and Durand, 2012; Martin et al., 2013). Interestingly, this could be caused by lack of inhibitory input to MNs (Fogarty et al., 2016). SOD1 spinal MNs remain larger than WT MNs through adulthood (Dukkipati et al., 2018). In contrast to the spinal MNs at this age, we found that SOD1 inhibitory interneurons in the ventral-most region had smaller soma size and yet had expanded surface area and volume of primary neurites compared to WT glycinergic interneurons. This is reminiscent of changes in corticospinal neurons: presymptomatic corticospinal neurons also have increased arborization but with smaller soma diameters (Ozdinler et al., 2011; Saba et al., 2016). In both sporadic and familial ALS patients, postmortem tissue shows degeneration of apical neurites in corticospinal neurons and smaller soma size (Genç et al., 2017). In SOD1 spinal interneurons in the present study, increased neurite surface area and volume of primary neurites were most prominent in the ventral interneurons in the RC region. This region also showed increased total neurite length and based on reconstructed, unpatched neurons, smaller soma volume. Altered neuronal size / arborization could alter the impact of other stressors (e.g., proteostasis, metabolic deficits, intracellular transport).

### After spike afterhyperpolarization

The AHP can contribute to a neuron’s firing rate such that a neuron with a large, long-lasting AHP will typically fire at a slower rate (typical of slow/type I MNs) than a neuron with small, fast-decaying AHP (typical of fast/type II MNs). In ALS patients, the AHP is shortened in MNs of patients with little force deficits and later elongated in patients with large force deficits (Piotrkiewicz and Hausmanowa-Petrusewicz, 2011), which could reflect early changes in physiology of vulnerable fast MNs and later, the remaining, less-vulnerable population of slow MNs. Similarly, SOD1 mouse MNs show shorter AHPs early, and longer AHPs around symptom onset (Quinlan et al., 2011; Jensen et al., 2020). Here we show that AHP is modestly shorter in duration in glycinergic SOD1 interneurons very early in life. In the regional analysis, AHP parameters were not found to be significantly altered in any of the 3 regions, but it is possible this was due to a loss in statistical power in the smaller groups. We did not find concomitant changes in minimum or maximum firing rates in GlyT2 interneurons, so the physiological importance of the altered AHP duration in interneurons is unclear.

### Inhibitory interneurons: Subtype specific effects

The interneurons in this study were heterogenous, and based on their location and electrophysiology, they likely were composed of V1 and V2b interneurons including commissural interneurons (V2b interneurons), RCs, and Ia inhibitory interneurons (both V1 interneurons), and perhaps inhibitory propriospinal interneurons (Willis and Willis, 1964; Restrepo et al., 2009; Bikoff et al., 2016; Flynn et al., 2017). Because of this heterogeneity, interneurons were separated into regions for further analysis, to examine more specifically which inhibitory interneurons were perturbed. Based on our regional analysis, interneurons in the ventral-most 100 μm from the white matter were altered the most in both excitability and morphology, with the most strongly depolarized PICs and larger primary neurites, more so than glycinergic interneurons within lamina IX and intermediate lamina VI and VII. Interneurons located in the region within 100 μm from the ventral white matter likely contain more RCs based on the location (Siembab et al., 2010) and electrophysiological characteristics (Perry et al., 2015; Bikoff et al., 2016). Glycinergic interneurons located in lamina IX are reminiscent of the locomotor-related GABAergic interneurons characterized by Nishimaru and colleagues, which were located within the ventrolateral spinal cord, surrounded MNs and were active during locomotor activity (Nishimaru et al., 2011). Indeed, these may be the same neurons: colocalization of GABA and glycine is entirely possible (Jonas et al., 1998; Allain et al., 2006). If the lamina IX interneurons are indeed locomotor-related, even the modest dampening of excitability observed here in SOD1 mice could contribute to changes in locomotor patterning as has been described in other studies (Quinlan et al., 2017; Allodi et al., 2021). Further studies are warranted to examine how inhibitory interneurons may be differently modulated in a subtype-specific manner in animal models of ALS.

### Functional implications for impaired inhibitory circuits

Outward signs of improperly functioning inhibitory interneurons may be apparent from subtle changes in motor patterning and could potentially be used as a biomarker of early ALS. Locomotor disturbances have been demonstrated *in vivo* in presymptomatic SOD1 mice during walking/running (Vinsant et al., 2013; Akay, 2014; Quinlan et al., 2017; Allodi et al., 2021). In the embryonic and neonatal spinal cord *in vitro*, increased duration of depolarizing events in MNs and a slower period of locomotor-related bursting has been described (Medelin et al., 2016; Branchereau et al., 2019), which indicate that even during development the functioning of locomotor circuits is already impaired. In presymptomatic adult mice capable of running on a treadmill, subtle locomotor differences are observed including advanced intermuscular phasing and a slower speed (Quinlan et al., 2017; Allodi et al., 2021) which indicate that before any large-scale neurodegeneration occurs, there are changes in patterns of activity in MNs and interneurons that could be exploited for use as a biomarker of ALS. At initial symptom onset, the first failure of the tibialis anterior manifests as abnormality in the swing phase of the ankle (Akay, 2014). Interestingly, silencing different interneuron populations (including inhibitory interneurons and cholinergic interneurons) reveals the contribution of each population: silencing cholinergic interneurons increases locomotor deficits in SOD1 mice (Landoni et al., 2019), while silencing inhibitory interneurons has no effect suggesting these interneurons are already impaired (Allodi et al., 2021).

### Clinical-translational implications

Evidence of MN hyperexcitability in ALS patients is present alongside signs of dysfunction in inhibition (Rothstein et al., 1992; Plaitakis and Constantakakis, 1993; Raynor and Shefner, 1994; Shefner and Logigian, 1998; Andreadou et al., 2008; Bae et al., 2013). Perhaps reducing excitability (with riluzole, for example) only results in a modest increase in lifespan (Bensimon et al., 1994) because it could be helpful to MNs while exacerbating inhibitory dysfunction by further depressing excitability of inhibitory interneurons like RCs. Thus, a more targeted approach to reduce MN excitability could be beneficial early in disease pathology. Stimulation of inhibitory nerves or certain protocols of transcranial stimulation could be explored, along with augmentation of inhibitory neurotransmission.

## Data Availability Statement

All data presented in this article were included in the Figures and Tables. Data were excluded only based on the criteria stated in the methods.

## Author contributions

CFC: acquisition of images and 3D reconstruction data, analysis and statistics, writing the manuscript

PRS: analysis of electrophysiology and statistical data, and revising the manuscript

EJR: 3D reconstructions and analysis, revisions to the manuscript

LTG: statistical analysis of electrophysiology and morphology, revisions to the manuscript

LMM: analysis of statistical data, revising the manuscript

KJL: analysis of 3D reconstruction data, revising the manuscript

ACP: analysis of electrophysiology data, revising the manuscript

MM: statistical analysis and effect size analysis, revisions to the manuscript

NK: conception and design of the work, revising the manuscript

KAQ: conception and design of the work, acquisition of electrophysiology and imaging data, analysis of electrophysiological data, statistics and writing the manuscript

All authors approved the final version of the manuscript; agreed to be accountable for all aspects of the work in ensuring that questions related to the accuracy or integrity of any part of the work are appropriately investigated and resolved; and all persons designated as authors qualify for authorship, and all those who qualify for authorship are listed.

## Acknowledgments

The authors made use of equipment supported by the Institutional Development Award (IDeA) Network for Biomedical Research Excellence from the National Institute of General Medical Sciences of the National Institutes of Health under grant number P20GM103430. The authors thank Drs. Ole Kiehn and Hans Zeilhofer for providing access and permission (respectively) to GlyT2eGFP mice for this study. The authors thank Claudia Fallini for comments on the initial drafts of the manuscript.

